# Potential universal PCR method to detect decapod hepanhamaparvovirus (DHPV) in crustaceans

**DOI:** 10.1101/2020.09.01.278721

**Authors:** Jiraporn Srisala, Dararat Thaiue, Piyachat Sanguanrut, Diva January Aldama-Cano, Timothy W. Flegel, Kallaya Sritunyalucksana

## Abstract

Parvoviruses that infect the hepatopancreas (HP) of the penaeid shrimp *Penaeus chinensis*, *P. monodon*, and *P. merquiensis* were previously called hepatopancreatic parvoviruses (HPV). They are now classified in the family *Parvoviridae*, sub-family *Hamaparvovirinae* as members of the same genus called *Hepanhamaparvovirus* and referred to as decapod hepanhamaparvovirus, designated here as DHPV. However, a virus that causes similar lesions in the HP of the giant river prawn *Macrobrachium rosenbergii* resembles hepanhamaparvoviruses by microscopy and histochemistry. Unfortunately, no genome information is yet available and PCR detection methods that work for DHPV in *P. monodon* do not work with *M. rosenbergii*. For hatchery samples of *M. rosenbergii* in Thailand with DHPV-like lesions, we hypothesized it might be possible to design primer pairs from 8 full DHPV genome sequences at GenBank for use in PCR detection of DHPV in *M. rosenbergii*. Using this strategy, we successfully designed a new set of primers and a PCR protocol called the DHPV-U method that gave an amplicon with DNA extracts from larvae of M. rosenberigii samples that showed DHPV-like lesions, while extracts from normal larvae gave none. DNA extracts from *P. monodon* infected with DHPV also gave amplicons. At the same time, the normal PCR method for DHPV in *P. monodon* gave no amplicon with the *M. rosenbergii* DNA extracts. The DHPV-U amplicons from *P. monodon* and *M. rosenbergii* shared 99% sequence identity, and *in situ* hybridization (ISH) assays using the DIG-labeled amplicon gave positive histochemical results in the HP tissue of both *P. monodon* and *M. rosenbergii*. The DHPV-U method is now being used in Thailand for detection of DHPV in both *P. monodon* and *M. rosenbergii*. Overall, the results support the proposal that the HP virus in *M. rosenbergii* is also a *hepanhamaparvovirus*. Based on 100% sequence identity of the target region in the currently published DHPV sequences at GenBank, the DHPV-U method may also work for detection of other DHPV isolates.

## 1. INTRODUCTION

Parvoviruses that infect the hepatopancreas (HP) of the penaeid shrimp *P. chinensis*, *P. monodon*, and *P. merquiensis* were previously called hepatopancreatic parvoviruses (HPV) (Lightner, 1996a). They are now classified in the family *Parvoviridae*, subfamily *Hamaparvovirinae* as members of a single species called decapod hepanhammaparvovirus 1 (Pénzes, et al., 2020), abbreviated here as DHPV. There were 8 full sequences listed at GenBank as *Hepandensovirus* and derived from the three penaeid shrimp species above. However, there is another virus that causes similar lesions in the HP of the giant river prawn *Macrobrachium rosenbergii* (Anderson, et al., 1990; Lightner, et al., 1994). Preliminary work by electron microscopy and histochemistry (Gangnonngiw, et al., 2009) suggested that the *M. rosenbergii* virus was also a parvovirus. However, no genome information was available, and it could not be detected using PCR methods designed for DHPV detection in *P. monodon* (Gangnonngiw, et al., 2009). DHPV was first reported from Thailand in *P. monodon* specimens (Flegel, et al., 1992) and it has been associated with retarded growth, disease and mortality in juvenile shrimp (Flegel, et al., 1999; Lightner, et al., 1993). However, DHPV (called HPV at the time) was later removed from the OIE list of reportable diseases after analysis showed there was no negative economic effect on the aquaculture industry due to the ability to exclude it from production facilities (Thitamadee, et al., 2016). Although HP lesions similar to those of DHPV in *P. monodon*, *P. megquiensis* and *P. chinensis* have been reported in other wild and cultured penaeid shrimp species including *P. esculentus*, *P. japonicus*, *P. semisulcatus*, *P. indicus*, *P. penicillatus*, *P. schmitti*, *P. vannamei* and *P. stylirostris* and in the Palaemonid shrimp *Macrobrachium rosenbergii* (Lightner, 1996b) genome sequence information is lacking, except for a few species (Walsh, et al., 2017), or it is insufficient to determine whether or how closely related they are to DHPV in *P. monodon*.

The most common diagnostic method that has been used for DHPV detection is histological analysis of the hepatopancreatic tubule cells by H&E staining to reveal pathognomonic lesions characterized by eosinophilic to basophilic, intranuclear inclusions contained in hypertrophied nuclei of hepatopancreatic tubule epithelial cells (Flegel, et al., 1999; Lightner, et al., 1993). Molecular detection can also be carried out by PCR following OIE standard methods (OIE, 2007) or by using DNA probes for *in situ* hybridization (Flegel, et al., 1999; Manjanaik, et al., 2005; Phromjai, et al., 2001). However, sensitivity of the methods may vary depending on the host species and/or its geographical location due to differences in some portions of the genomes among the DHPV isolates (Dhar, et al., 2014; Phromjai, et al., 2001; Tang, et al., 2008).

From 2016-2018, a project to develop specific pathogen free (SPF) *M. rosenbergii* contacted our research unit to diagnose the cause of unexpected mortality in larvae from a screening program to select a founder stock. The affected larvae showed typical DHPV-like lesions, as had previously been reported (Anderson, et al., 1990; Gangnonngiw, et al., 2009). At the same time, several full sequences of DHPV had accumulated at GenBank, and we hypothesized that, if the virus in *M. rosenbergii* was also a *hepanhamaparvovirus*, we might be able to design a PCR detection method for it from regions of high sequence identity among the reported isolates of DHPV. Here we report the success of this approach and its confirmation by *in situ* hybridization analysis. At the same time, analysis of the amplicon sequence supports the proposal that the lesions in *M. rosenbergii* are also caused either directly or indirectly by a *hepanhamaparvovirus*.

## 2. MATERIALS AND METHODS

### 2.1. Shrimp specimens

Two batches of PLs of *Macrobrachium rosenbergii* (~7-14 mm in length) exhibiting signs of a suspected disease outbreak were obtained in December 2017 and January 2018 from a hatchery in Suphanburi province, Thailand. Batch #1 (40 PLs 12-14 mm in length) was divided into 2 subgroups: one subgroup (30 PLs) was divided into 3 tubes containing 10 PLs each in 500 μl of TF lysis buffer (50 mM Tris-HCl, 100 mM EDTA, 50 mM NaCl, 2% SDS, 10 μg/ml proteinase K, pH 9.0) for DNA extraction, while the remaining 10 were fixed in Davidson’s fixative for histological analysis individually in 10 paraffin blocks. Similarly, Batch #2 (114 PLs 7-10 mm in length) was divided into 2 subgroups: one subgroup (100 PLs) was divided into 10 tubes containing 10 PLs each in 500 μl of TF lysis buffer as above while the remaining 14 PLs were fixed with the Davidson’s fixative. The fixed PLs were processed for histological analysis in two paraffin blocks containing 7 PLs each. Archived DNA extracts and an archived paraffin block of tissue from *P. monodon* infected with DHPV1 were used for analysis in comparison with the samples from *M. rosenbergii*.

At the time this work was carried out, there was no official standard of the Ethical Principles and Guidelines for the Use of Animals of the National Research Council of Thailand (1999) for invertebrates. However, its principles for vertebrates were adapted for prawn specimens. The guidelines of the New South Wales (Australia) state government for the humane harvesting of fish and crustaceans were followed (http://www.dpi.nsw.gov.au/agriculture/livestock/animal-welfare/general/fish/shellfish) with respect to processing of the prawns for analysis. The saltwater/ice slurry method was used as recommended in the Australian guidelines.

### 2.2 Histological analysis

For histological analysis, the living PL of *M. rosenbergii* specimens were stunned an ice slurry and immediately fixed whole with Davidson’s fixative solution overnight before processing for hematoxylin and eosin (H&E) staining (Bell Lightner, 1988). After that, the hepatopancreatic tissue of each specimen was screened by light microscopy for the presence of DHPV-like lesions. Sections of the same paraffin-embedded tissues were used for *in situ* hybridization assays.

### 2.3 Nucleic acid extraction for PCR amplification

The 13 *M. rosenbergii* subsamples of 10 PL each were processed first by removal of eyestalks (to avoid PCR interference) before homogenization in 500 μl of TF lysis buffer [50 mM Tris-HCl (pH 9.0), 100 mM EDTA, 50 mM NaCl, 2%SDS, 10 μg/ml Proteinase K] and incubated for 1 h at 60-65°C. Total DNA was purified following the standard phenol: chloroform: isoamyl alcohol protocol (Sambrook Russell, 2001) and the DNA pellet obtained was resuspended with 30 μl of DNase/RNase free water. Concentration of DNA was determined using a dsDNA BR assays on a Qubit 3.0 Fluorometer (Life Technologies) and stored at −20°C until used.

### 2.4 Design of PCR primers

A new set of primers for DHPV detection was designed base on highly conserved sequences of 8 selected DHPV genome sequences available at the GenBank database (**Table 1**). The name assigned to this primer set was decapod *hepanhamaparvovirus* universal primers or DHPV-U primers. The nucleotide sequences of the DHPV-U primers and the existing OIE primers used in the study are listed in Table 2.

**Table 1.** Nucleotide sequences of complete or almost complete genomes used for multiple sequence alignment analysis for the construction of the DHPV-U primers. The virus names used in the table are those that were used at GenBank at the time of writing. In this article, the penaeid shrimp binomials used are according to Holthuis (1980) and Flegel (2007).

| Host Species | Virus name | Accession No. |
| --- | --- | --- |
| *P. monodon* | Penaeus monodon hepandensovirus 1 | DQ002873.1 |
|  | Penaeus monodon hepandensovirus 2 | JN082231.1, |
|  | Penaeus monodon hepandensovirus 3 | EU588991.1 |
|  | Penaeus monodon hepandensovirus 4 | FJ410797.2 |
| *P. chinensis* | Penaeus chinensis hepandensovirus | GU371276.1, AY008257.2, EU247528.1 |
| *P. merquiensis* | Penaeus merguiensis hepandensovirus | DQ458781.4 |

**Table 2.**
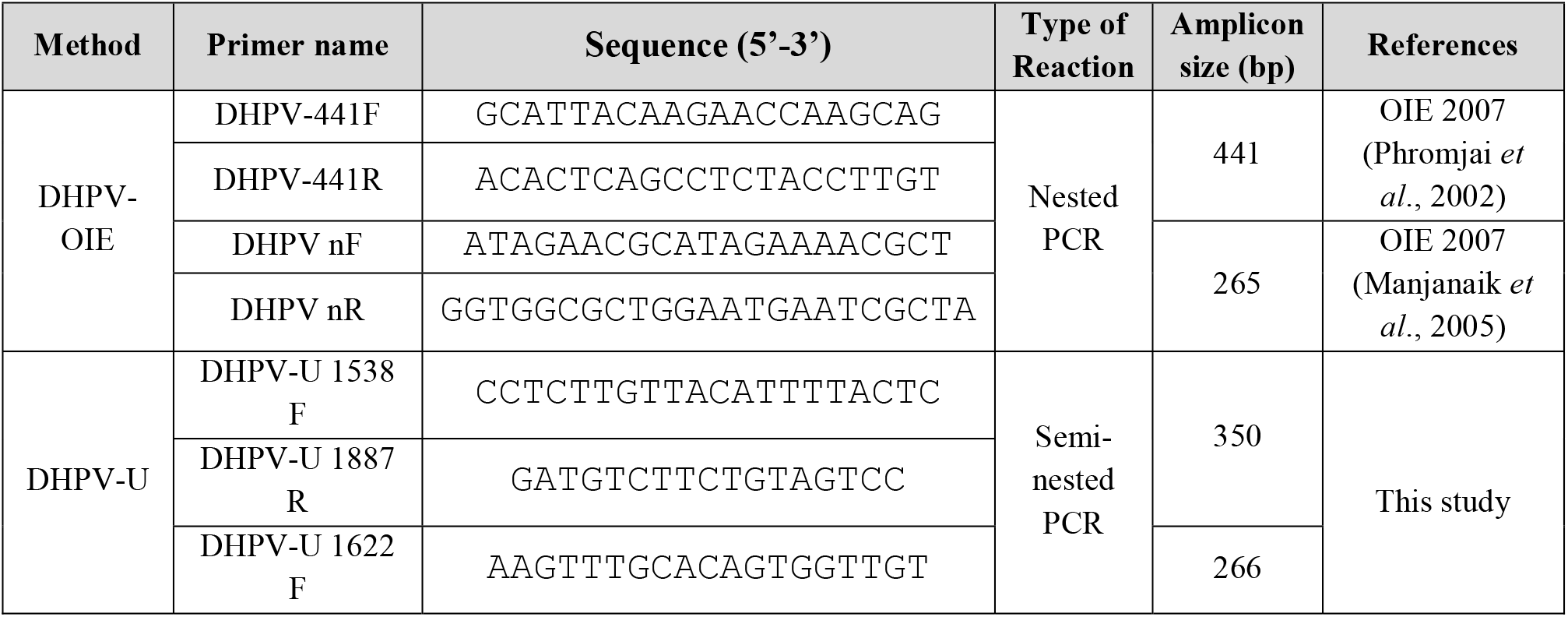
Primers used for PCR amplification in this study.

### 2.5 PCR methods

For the DHPV-OIE method, the primers listed in Table 2 by Phromjai *et al* (Phromjai, et al., 2002)and Manjanaik *et al* (Manjanaik, et al., 2005) were used, and the reaction was performed following the OIE-manual PCR detection method. For the DHPV-U method, the first step PCR reaction was performed in a 12.5 μl mixture containing 1X OneTaq Hot Start Master Mix (NEB), 0.4 μM of each DHPV-U 1538F and DHPV-U 1887R primer and 20 ng of DNA template. The PCR protocol was initial denaturation for 5 min at 94°C followed by 30 cycles of denaturation for 30 s at 94°C, annealing for 30 s at 55°C and extension for 45 s at 72°C with a final extension for 5 min at 72°C. For the semi-nested PCR step, the 12.5 μl reaction mixture contained 1X OneTaq Hot Start Master Mix (NEB), 0.2 μM of each DHPV-U 1622F and DHPV-U 1887R primers and l μl of PCR product from the first-step PCR reaction. The PCR protocol was initial denaturation for 5 min at 94°C followed by 25 cycles of denaturation for 30 s at 94°C, annealing for 30 s at 55°C and extension for 30 s at 72°C with a final extension for 5 min at 72°C. The amplicons were analyzed by 1.5% agarose gel electrophoresis with ethidium bromide staining and using a DNA ladder marker (2 log DNA ladder from New England Biolabs, USA). Amplicon bands were observed under UV light. The expected amplicons were for light infections one 266 bp band and for heavy infections one 266 bp band plus one 350 bp band. The PCR products obtained were cloned into pGEM-T-Easy vector and subjected to bi-directional sequencing (Macrogen, Korea).

### 2.6 Specificity and sensitivity of the DHPV-U method

To determine the sensitivity of the DHPV-U method, a recombinant pGEM-T plasmid was constructed to contain a DHPV amplicon and it was used as a template at 10-fold serial dilutions in corresponding PCR reactions. The highest dilution that still gave a visible band on the agarose gel was considered the lowest detectable quantity of target DNA, and the equivalent copy number was calculated using Avogadro’s number against the molar quantity of plasmid DNA. To determine the specificity of the DHPV-U primer, crustacean samples severely infected with other viruses were tested. These included archived DNA and RNA extracted from *P. vannamei*] severely infected with either white spot syndrome virus (WSSV), yellow head virus (YHV) or Infectious myonecrosis virus (IMNV), and from *P. monodon* severely infected with infectious hypodermal and haematopoietic necrosis virus (IHHNV) or Laem Singh virus (LSNV). For WSSV, YHV and IMNV, the IQ2000 kits for detection (GeneReach, Taiwan) of each virus were used. For IHHNV, the levels of infection were determined by the OIE method following Tang et al. (Tang, et al., 2007). For LSNV, the PCR method followed Sritunyalucksana et al., 2006 (Sritunyalucksana, et al., 2006). For the YHV-, IMNV-, and LSNV-infected shrimp, the RNA was extracted and subjected to the Superscript™ III Reverse transcriptase (Invitrogen, USA) before the cDNA was used as the template for the DHPV-U method. Archived DNA extracts from *P. monodon* infected with DHPV were also used to compare the DHPV-O and DHPV-U methods.

### 2.7 *In situ* hybridization (ISH) assays using a DHPV-U-derived probe

The *in situ* hybridization tests were carried out using paraffin blocks containing DHPV-PCR positive PL of *M. rosenbergii* collected in this study and using archived blocks of DHPV-PCR positive juvenile stages of *P. monodon*. Digoxygenin (DIG)-labeled DNA probes (Roche, Germany) were generated by PCR according to the manufacturer’s instructions. The primers used for the labeling reactions were DHPV-U 1622F and DHPV-U 1887R. The PCR reaction was performed in 25 μl containing 0.4 μM of each primer, 1X PCR buffer [200 mM Tris-HCl (pH 8.4), 500 mM KCl], 1X PCR DIG labeling mix (Roche, USA), 1.5 mM MgCl_2_, 1.25 U Taq DNA polymerase (Invitrogen, USA) and 2×10^6^ copies of a plasmid clone containing the DHPV amplicon. The labeled PCR product was purified using a PCR purification kit (Geneaid, Taiwan) and stored in DNase/RNase free water at −20° C until used.

The protocol for ISH was as previously described (Sritunyalucksana, et al., 2006; Tangprasittipap, et al., 2013). Briefly, tissue sections were deparaffinized and rehydrated before being digested with 200 μl of 5 μg/ml Proteinase K (Invitrogen, USA) in TNE buffer for 1 h at 37°C. The sections were incubated in 0.5M EDTA at room temperature (~25°C) for 1 h before being fixed with ice cold 0.4% formaldehyde solution for 5 min and immersed in distilled water for 5 min. The sections were equilibrated with pre-hybridization solution [4 × SSC and 50% (v/v) deionized formamide] at 37°C for 1 h. After that, the sections were replaced with hybridization solution containing the DIG-labeled probe (approximately 400 ng/ slide) and covered with a coverslip. The control reaction without probe was included in a separate container. The hybridization reaction was incubated at 42°C overnight in a humid chamber. After incubation, the sections were washed sequentially for 10 min with 2X SSC, 15 min with 2XSSC at 37°C, 15 min with 1XSSC at 42°C, 15 min with 0.5X SSC at 42°C and 5 min with buffer I [100 mM Tris–HCl and 150 mM NaCl, pH 7.5] at room temperature. After washing, the sections were equilibrated with buffer II [Buffer I containing 0.5% Blocking reagent (Roche, Germany)] at room temperature for 1 h before incubation with alkaline phosphatase-conjugated anti-digoxigenin antibody (1:500 dilution). The sections were washed 2×10 min with buffer I and equilibrated in detection buffer (100 mM Tris– HCl and 100 mM NaCl, pH 9.5). The signal was developed by addition NBT-BCIP substrate (Roche, Germany) in the dark and counterstaining was accomplished with Bismarck brown Y (Sigma, USA). The slides were observed and photographed using an Olympus microscope with a digital camera.

## 3. RESULTS AND DISCUSSION

### 3.1. A detection method for DHPV established using DHPV-U primers

A total of 8 complete or nearly complete genomes of DHPV derived from *P. chinensis* (3), *P. merquiensis* (1) and *P. monodon* (4) were selected from the GenBank database on 09 September 2018. There were also records for an additional 16 sequences, two of which were redundant to 2 of the 8 selected isolates and 14 that were associated with retracted reports and included a record from *P. indicus*. These sequences are not included in the analysis shown in **Fig. 1**. Multiple sequence alignment of the 8 selected sequences revealed that sequence identity in many regions were highly conserved at 100% identity (**Fig. 1** and **Supplementary Fig. 1**). Primers for detection of DHPV were designed from one such region 1538-1556 bp (green), 1622-2640 (blue) and 1871-1887 (pink), as shown in **Fig. 1A**. This region also showed 100% sequence identity in all 14 of the other GenBank sequences not included in the analysis shown in Fig. 1. Theoretically, based on 100% identity in the target sequence across all the shrimp hosts, the primers should be effective with all 8 of the selected viral types and might also give amplicons with closely related but currently unknown isolates from other species or geographical regions. The locations of the DHPV-U primers in comparison with those used for the OIE method are shown in Figure 1B using the whole genome sequence of DHPV accession no DQ002873.1.

**Figure 1.**
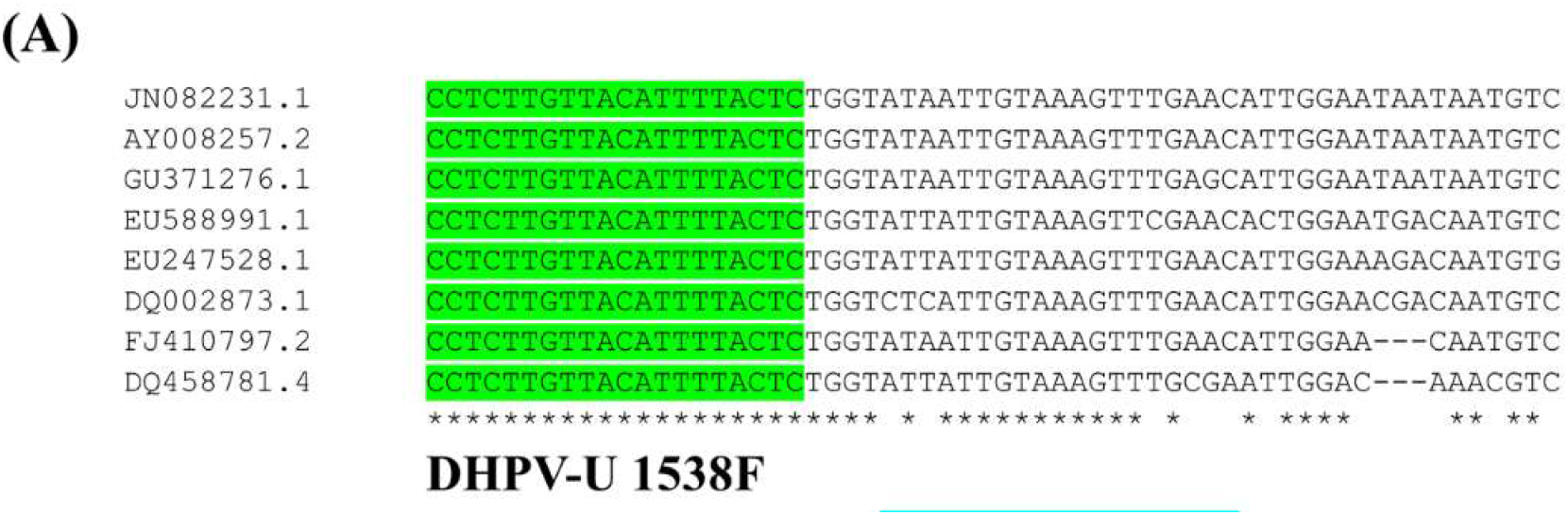
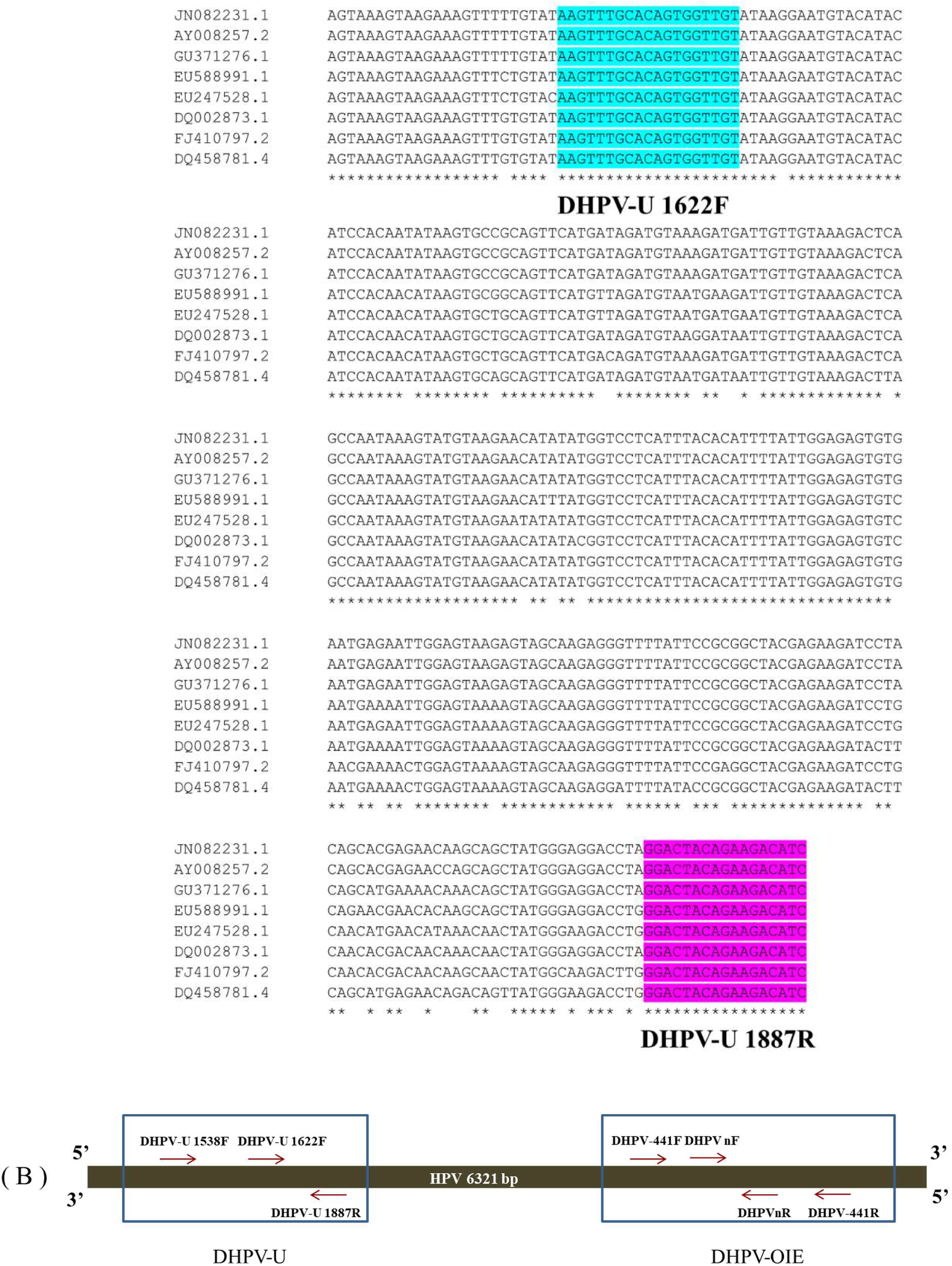
(A) Multiple sequences alignment of 8 DHPV sequences available at the NCBI database, as listed in Table 1. The conserved nucleotides were indicated by asterisks (*). The highlighted regions were used to design DHV-U primers; DHPV-U 1538F (green), DHV-U 1622F (blue) and DHV-U 1887R (pink). the DHV-U primer sequences and lack of cross reactions with other important viral pathogens of crustaceans. (B) Graphical Primer design from the DHPV-OIE method compared to the DHPV-U method presented in this study. The DHPV sequence belongs to NCBI accession no. DQ002873.1. Red arrows indicate the forward and reverse primer positions.

### 3.2. Specificity and sensitivity testing of the DHPV-U method

Using the semi-nested DHPV-U PCR protocol with DNA or cDNA templates derived from penaeid shrimp infected with the viruses WSSV, YHV, IMNV, IHHNV and LSNV gave no amplicons (**Fig. 2**). In contrast, the DNA template from *M. rosenbergii* infected with DHPV gave a positive test result using the DHPV-U PCR protocol (lane 6). The results revealed no cross reactivity of DHPV-U primers with other common shrimp viruses (lanes 1-5).

**Figure 2.**
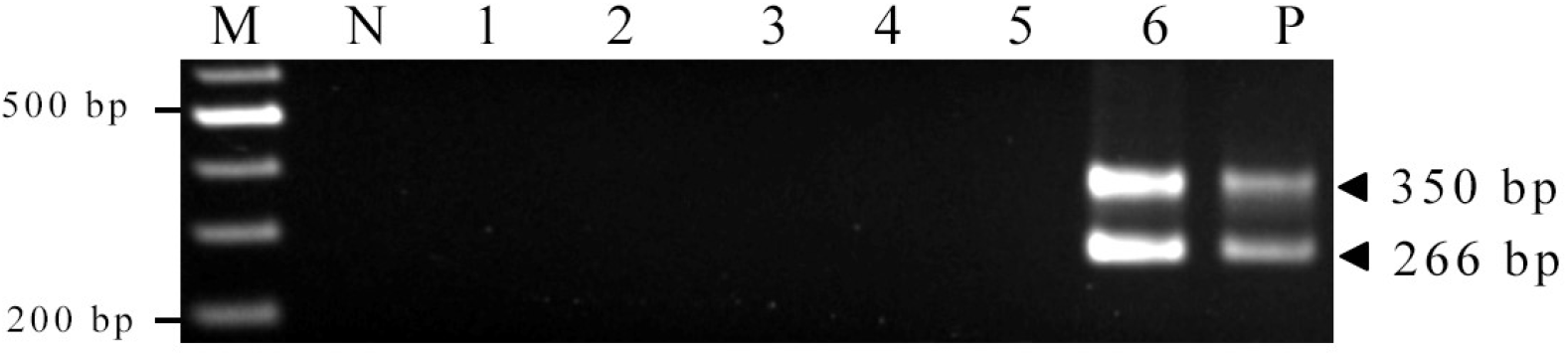
Specificity testing for the DHPV-U PCR detection method. Agarose gel electrophoresis analyses of the DHPV-U reaction solutions by DNA or cDNA templates from YHV-infected shrimp (lane 1), LSNV-infected shrimp (lane 2), IMNV-infected shrimp (lane 3), WSSV-infected shrimp (lane 4), IHHNV-infected shrimp (lane 5), and DHPV-infected prawn (lane 6). M = 2 log DNA marker, N = Negative Control, P = Positive control. The expected sizes of PCR products amplified by DHPV-U PCR were 266 and 350 bp.

Testing the sensitivity of the DHPV-U method using a serially diluted plasmid template containing a DHPV target (0 – 2×10^5^ plasmid copies per reaction tube) (**Fig. 3**), revealed that the semi-nested DHPV-U method could detect as little as 2 copies/reaction. A second test was carried out using templates that contained both DHPV-free shrimp DNA (20 ng/reaction) plus DHPV-U plasmid preparation from 0-200 plasmids/reaction. The results (**Fig. 3B**) show that the lowest copy number of DHPV-U plasmid that could be detected by the DHPV-U method was 50 copies when 20 ng of host DNA was included in the reaction mix. This revealed a strong negative influence of the host DNA on the sensitivity of the DHPV-U method. This effect has been studied and demonstrated in several models (Cogswell et al., 1996; Handschur et al., 2009). Cogswell et al. (1996) demonstrated that host DNA can interfere with specific DNA amplification for *Borrelia burgdorferi* by PCR and may even lead to false negative results. The sensitivity of pathogen detection is also reduced in next generation sequencing in specimens containing high background human DNA. Several methods have been proposed to solve this problem, including dilution of the template, concentration and extensive washing of the DNA template.

**Figure 3.**
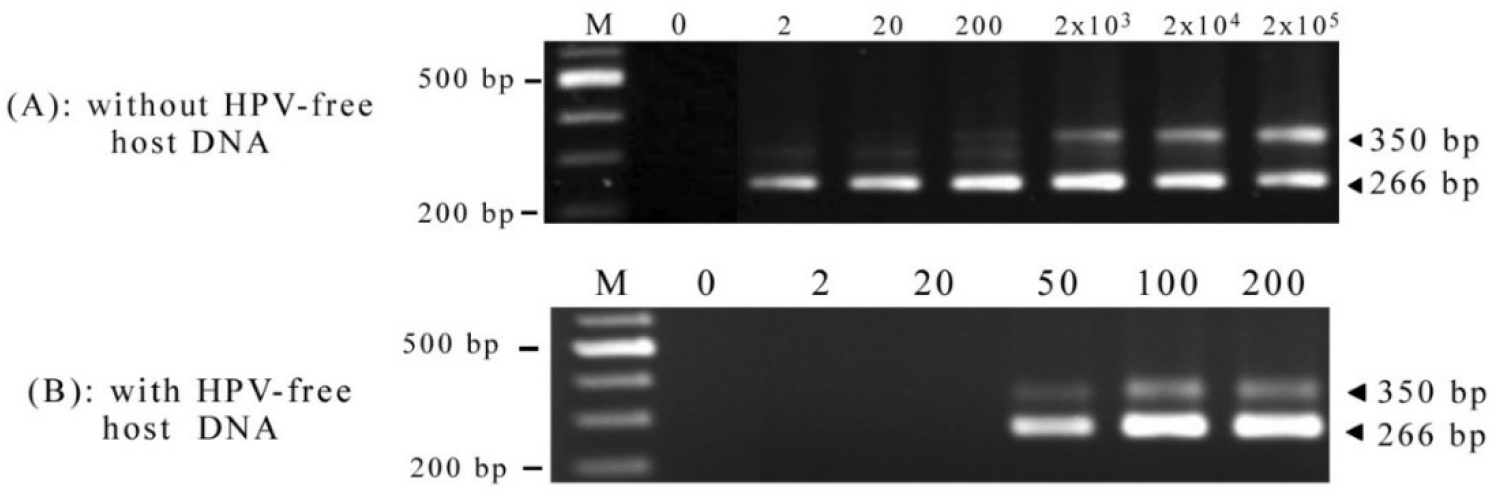
Sensitivity testing for the DHPV-U method. (A) Agarose gel electrophoresis analyses the DHPV-U amplicons from PCR using serially diluted DHPV-U plasmid templates at 0 to 2×10^5^ plasmids/reaction. (B) Agarose gel showing the effect of host DNA addition (20 ng/reaction) to reactions containing DHPV-U plasmid from 0-200 copies.

The method compares favorably with the sensitivity of the previously published OIE one-step PCR method (OIE, 2007) and the previously published nested PCR method (Manjanaik et al., 2005) for DHPV in *P. monodon*.

### 3.3 Histological analysis of DHPV-infected *M. rosenbergii*

To confirm the positive PCR results, histological analysis of the affected HP tissue was necessary to confirm the diagnosis since there were no specific gross signs to detect the presence DHPV. Histological examination of tissue sections from the fixed samples of *M. rosenbergii* PL revealed the presence of spherical to ovoid intranuclear inclusions in the HP tubule epithelial cells of some of the specimens. From the first Batch of 10 specimens, only 1 specimen showed typical DHPV-like lesions in the HP. From Batch 2 with 2 slides each with 7 specimens each (total 14), 6 showed DHPV like lesions. An example is shown in Fig 4. These results were sufficient to confirm the PCR results from the 2 batches of shrimp.

**Figure 4.**
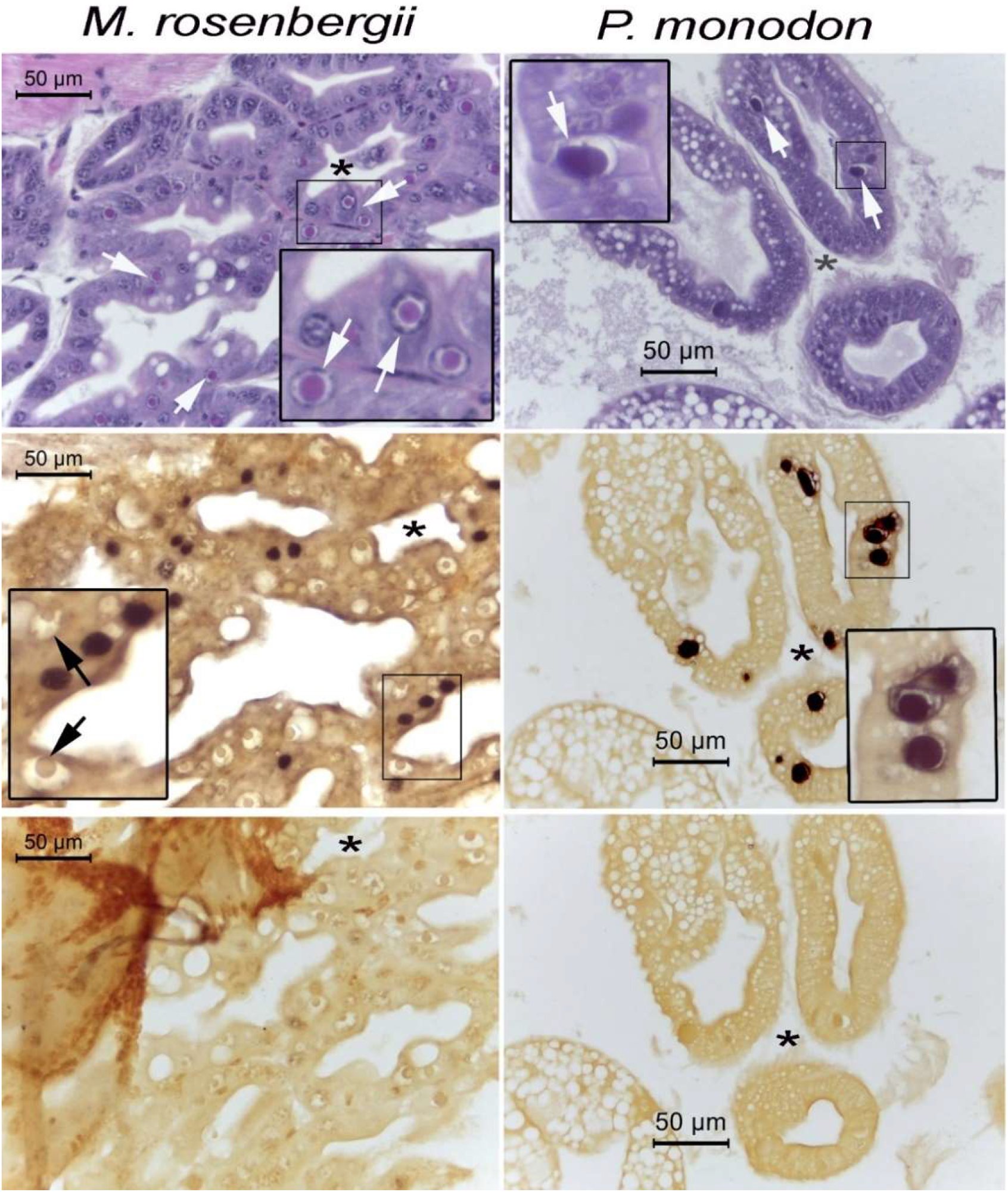
Example photomicrographs of histopathology and ISH reactions with HP tissue of *M. rosenbergii* and *P. monodon*. **Row 1**. H&E staining showing DHPV-like intranuclear inclusions in tubule epithelial cells marked with white arrows. Inserts show magnified regions. **Row 2**. Positive ISH reactions (black staining) in locations matching the regions of the intranuclear inclusions in the adjacent sections in Row 1. **Row 3**. Negative controls for the ISH reactions (no probe). Asterisks in the adjacent tissue sections indicate that same relative position for the photomicrographs in each column.

The intranuclear inclusions varied in size but were all eosinophilic, characteristic of early stage DHPV lesions (**Figure 4**, **row 1**, white arrows). These were similar to the suspected DHPV-like lesions previously observed in samples of *M. rosenbergii* PL and broodstock reported by (Gangnonngiw, et al., 2009). However, in the broodstock, some of the larger inclusions stained basophilic. The fixed hepatopancreas of a *P. monodon* specimens known to be infected with DHPV were used to compare lesions of similar morphology and staining characteristics in its HP tubule epithelial cells. Example photomicrographs are shown in **Fig. 4** (row 1, white arrows).

### 3.4 *In situ* hybridization confirmed DHV infections in *M. rosenbergii*

To confirm that the DHPV-like histological lesions in the HP of the *M. rosenbergii* were associated with the positive DHPV-U PCR reactions, *in situ* hybridization assays were carried out using a DIG-labeled probe derived from a DHPV-U-PCR amplicon. Tissue sections from *M. rosenbergii* larvae from batches that tested positive using the DHPV-U PCR method and from *P. monodon* hepatopancreatic tissue known to be infected with DHPV both gave positive *in situ* hybridization reactions in the nuclei of HP tubule epithelial cells (dark staining against the brown counter-stain) using DIG-labeled probes for DHPV (**Fig. 4**, row 2). These reactions were at similar intensity and in the same tissue areas where the intranuclear inclusions were seen with hematoxylin and eosin (H&E)-stained, adjacent tissue sections (**Fig 4**, **row 1**). No reactions occurred in the control slides processed with no probe present (**Figs. 4**, **row 3**).

Curiously, the positive ISH reactions that occurred in the *M. rosenbergii* specimens did not arise from the intranuclear inclusions associated with DHPV infections but instead in nuclei without such inclusions present. In contrast, the ISH reactions in *P. monodon* did occur with DHPV inclusions. We have no explanation for this anomaly. We speculate that the inclusion structures or contents in *M. rosenbergii* may prevent hybridization with the probe in some unknown way, such that only nuclei undergoing genomic viral DNA synthesis prior to inclusion formation give positive ISH results. Our attempts to rectify the situation with additional proteinase K treatment or with sodium hydroxide treatment prior to the ISH reaction did not change the situation. This is curious because earlier work (Gangnonngiw, et al., 2009) showed by confocal laser microscopy that fluorescence from stained nucleic acid in the inclusions was lost or reduced by treatment with DNase 1 or with mungbean nuclease specific for single-stranded DNA, even though the inclusions remained intact. Thus, if the DNase enzyme could penetrate the structure of the inclusions, it is curious that the labeled nucleic acid probe apparently could not and/or was unable to hybridize with the viral DNA. Alternatively, it is possible that DHPV in *M. rosenbergii* is present in nuclei of normal histological appearance and that the distinctive, eosinophilic to basophilic inclusions arose from some direct or indirect associated cause, although to us this seems unlikely.

### 3.5 Comparison of DHPV-OIE and DHPV-U with field samples

These tests employed 13 DNA extracts from pooled PL samples (10 each) of *M. rosenbergii* that were suspected of being infected with DHPV and with 11 archived DNA extracts from *P. monodon* infected with DHPV1. When tested for DHPV using the OIE recommended method normally used for detection of DHPV in *P. monodon* (**Fig. 5A**), all 13 *M. rosenbergii* samples gave negative results. When the same DNA extracts were used as templates for the DHPV-U method 11/13 (85%) (**Fig. 5B**) gave positive test results. It is possible that the 2 samples of 10 that gave negative results did not contain even 1 PL infected with DHPV, since only 1 of the 10 PLs sampled for histological examination showed DHPV lesions. This indicated that DHPV was not highly prevalent in that batch of PLs such that an occasional sample of 10 might consist of uninfected individuals or might contain a lightly infected individual that yielded too little viral DNA to be detected after mixing with host DNA form 9 other uninfected individuals.

**Figure. 5.**
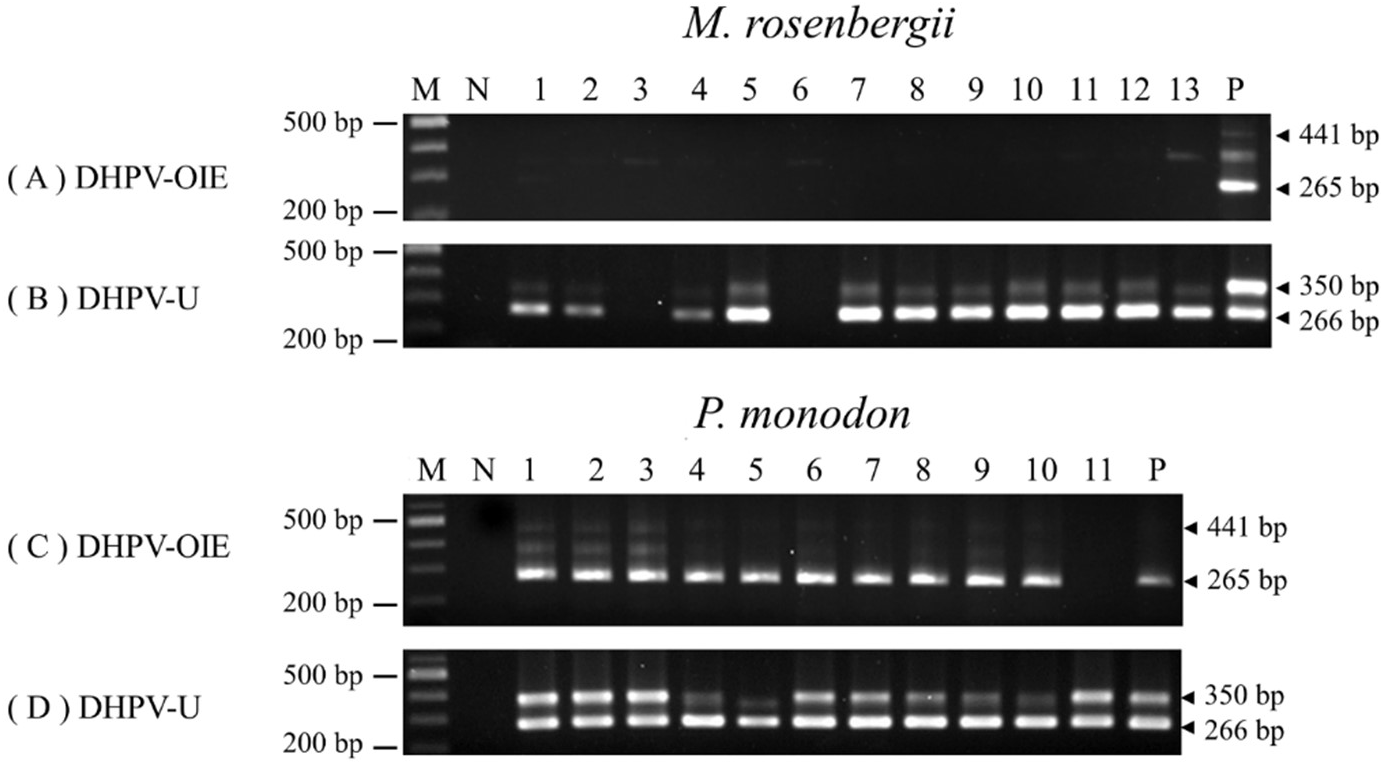
Comparison of DHPV detection in 13 *M. rosenbergii* and 11 *P. monodon* samples using the standard OIE detection method or the DHPV-U method. (A & B) Agarose gel results from using the 2 methods with *M. rosenbergii* samples and showing that the DHPV-OIE method does not work, while the DHPV-U method does. (C & D) Agarose gel results from using the 2 methods with *P. monodon* samples showing that both methods work with *P. monodon*. N = Negative control, M = 2 log DNA marker and P = Positive control.

When the DHPV-O and DHPV-U methods were used with 11 archived DNA extracts from *P. monodon* samples that showed DHPV lesions, 11/11 samples gave positive amplicons with the DHPV-O method although the band for sample 11 was very light (indicating a low level infection) and does not show up in photograph in Fig. 5C. The DPHV-U method also gave but 11/11 positive amplicons with the *P. monodon* samples. This clearly revealed that the DHPV-U method could be used for the different DHPV types in *P. monodon* and *M. rosenbergii*.

Next, 5 PCR amplicons (350 bp) from *M. rosenbergii* were arbitrarily selected and subjected to sequencing (Macrogen, Korea) and analysis. All 5 sequences were nearly identical, differing from one another by only 1 or 2 bases, always at different positions (Supplementary Fig. 1). These gave a consensus sequence that was used for an nBLAST search against the GenBank database. The top hit of 100% coverage and 99.4% identity (311/313 bases, excluding the primers) was for *P. monodon* hepandensovirus 1 (DQ002873.1) (**Fig. 6**).

**Figure 6.**
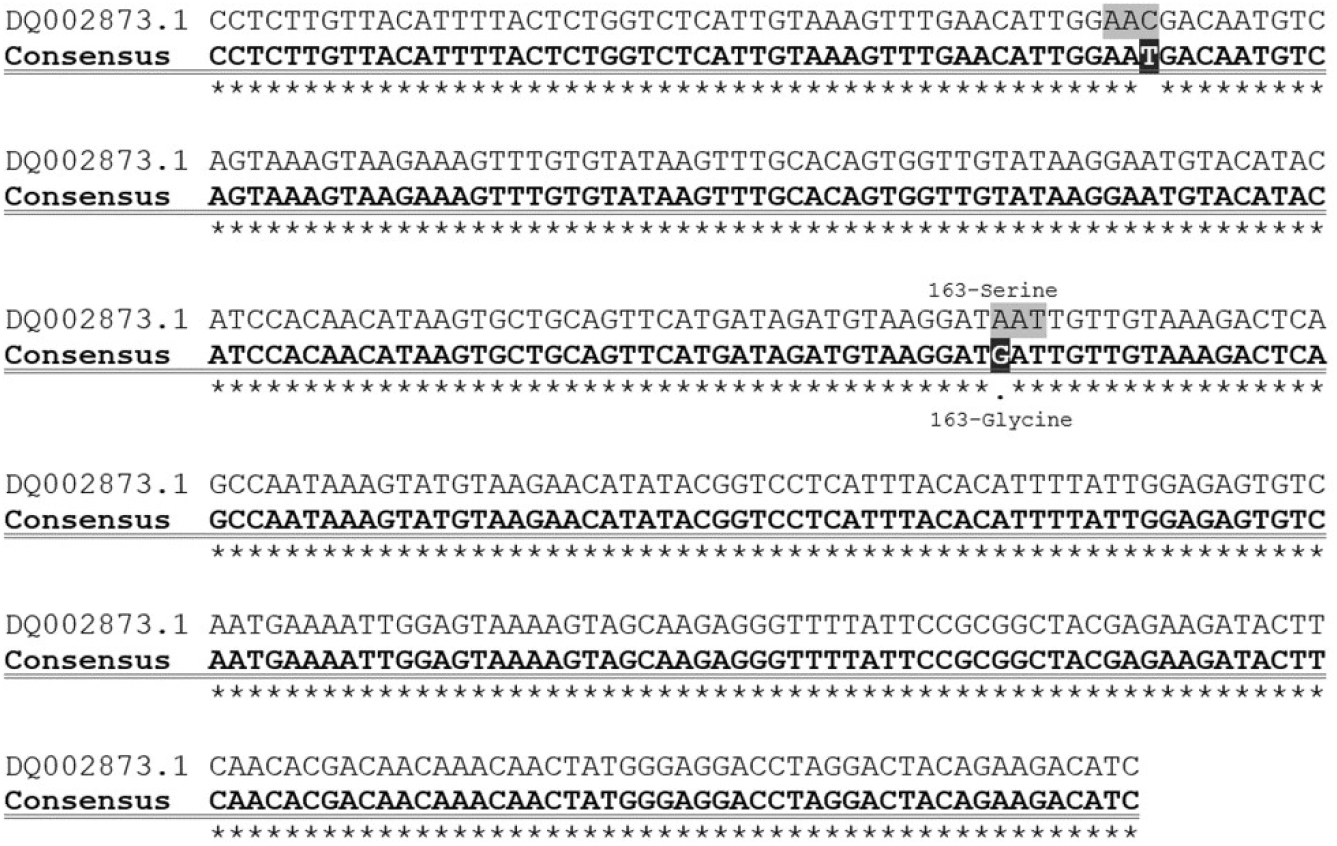
Sequence alignment obtained for the consensus sequence of 5 separate DHPV-U-PCR amplicon clones obtained from *M. rosenbergii* with the matching region of the Blast-n top-hit *Penaeus monodon* hepandensovirus 1 (DQ002873.1). There are two base differences giving a sequence identity of 348/350 = 99.4%. The first difference (position 48-51) is synonymous for serine while the second (position 163-165) is non-synonymous but is a semi-conserved change from serine to glycine.

Overall, the results revealed that the DHPV-U method could be used to screen for DHPV in both *P. monodon* and *M. rosenbergii* in Thailand without any negative consequences in terms of sensitivity or specificity. Indeed, it is somewhat more sensitive than the DHPV-O method at 50 copies when mixed with host DNA compare to 340 for the DHPV-O method (OIE, 2007). Some laboratories have already adopted the DHPV-U method since it provides some convenience when DHPV testing is being carried out with both species. On the other hand, the two methods together would be useful in determining whether the types of DHPV in *P. monodon* and *M. rosenbergii* are cross infective. This could now be determined easily in laboratory studies using the two methods to follow the infections.

## Conclusions

The DHPV-U method described herein can be used to screen for DHPV in both *M. rosenbergii* and *P. monodon*. Indeed, it is currently being applied for the screening of broodstock and larvae in a program aimed at developing an SPF stock of *M. rosenbergii* in Thailand. It is hoped that such a stock would ultimately provide shrimp farmers with PL free of relevant major pathogens. However, there is still interest in determining the full sequence of the DHPV type or types prevalent in the natural and imported sources of *M. rosenbergii* broodstock that are currently being used in Thailand. Given the high sequence conservation in existing GenBank records for the target sequence of the DHPV-U method, it may be useful in broad, preliminary screening for previously unknown isolates of DHPV in other fresh water, brackish water or marine animals.

## Acknowledgements

We would like to thank the financial support from the Royal Society International Collaboration Awards 2019, the Global Challenges Research Fund (GCRF) to Prof. Grant D. Stentiford (Cefas/UK) and Dr. Kallaya Sritunyalucksana (BIOTEC, NSTDA/Thailand).

